# *Tet2* loss suppress α-synuclein pathology by stimulating ciliogenesis

**DOI:** 10.1101/2024.08.02.606408

**Authors:** Emmanuel Quansah, Naman Vatsa, Elizabeth Ensink, Jaycie Brown, Tyce Cave, Miguel Aguileta, Emily Kuhn, Allison Lindquist, Carla Gilliland, Jennifer A. Steiner, Martha L. Escobar Galvis, Milda Milčiūtė, Michael Henderson, Patrik Brundin, Lena Brundin, Lee L. Marshall, Juozas Gordevicius

## Abstract

There are no approved treatments that slow Parkinson’s disease (PD) progression and therefore it is important to identify novel pathogenic mechanisms that can be targeted. Loss of the epigenetic marker, *Tet2* appears to have some beneficial effects in PD models, but the underlying mechanism of action is not well understood. We performed an unbiased transcriptomic analysis of cortical neurons isolated from patients with PD to identify dysregulated pathways and determine their potential contributions to the disease process. We discovered that genes associated with primary cilia, non-synaptic sensory and signaling organelles, are upregulated in both early and late PD patients. Enhancing ciliogenesis in primary cortical neurons via sonic hedgehog signaling suppressed the accumulation of α-synuclein pathology *in vitro*. Interestingly, deletion of *Tet2* in mice also enhanced the expression of primary cilia and sonic hedgehog signaling genes and rescued the accumulation of α-synuclein pathology and dopamine neuron degeneration *in vivo*. Our findings demonstrate the crucial role of *Tet2* loss in regulating ciliogenesis and potentially affecting the progression of PD pathology.

## INTRODUCTION

Parkinson’s disease (PD) is a common neurodegenerative disorder characterized by the progressive degeneration of dopamine neurons in the substantia nigra and the formation of Lewy bodies that contain aggregated α-synuclein (αSyn). In experimental PD models, αSyn pathology has been shown to spread throughout the brain to anatomically connected regions ^1,2^. Based on cross-sectional neuropathological studies, it has been suggested that αSyn aggregates spread in a similar fashion between brain regions in PD, and this progressive involvement of additional brain regions is thought to contribute to increased symptom severity and the emergence of new symptoms in PD ^3–5^. Both genetic and epigenetic factors can contribute to PD pathogenesis and may regulate αSyn aggregation and spread ^6,7^. Identifying genetic factors impacting the risk for the initial αSyn aggregation in circumscribed parts of the nervous system (prior to the manifestation of the disease-associated motor symptoms), as well as genes that influence the spread of αSyn pathology from these sites can reveal novel processes that can be targeted in attempts to slow PD progression.

The vast majority of PD cases are sporadic and heritable familial mutations account for about 5-15% of cases (depending on geographical regions and ethnicity). Mutations in genes such as *SNCA, GBA1, LRRK2, DJ-1*, and *PRKN* confer significant risk for PD pathogenesis. Ten-eleven translocation-2 (*TET2*) is another gene that has recently been linked to PD ^8,9^. Interestingly, DNA methylation abnormalities found in the *TET2* enhancer increases as Lewy pathology worsens ^8^, suggesting a possible correlation between TET2 and αSyn-rich Lewy pathology. However, very little is known about how TET2 activity influences αSyn pathology. TET2 belongs to the TET family of dioxygenases, which includes TET1, TET2 and TET3. They are evolutionarily conserved and are important for DNA methylome regulation ^10^. TET2 catalyzes the oxidation of 5-methylcytosine to 5-hydroxymethylcytosine (5hmC). 5hmC and its oxidized derivatives may eventually be replaced with an unmodified cytosine by base excision repair, resulting in demethylation ^11^. Emerging evidence suggests that the TET family is important for cellular development, differentiation, and reprogramming ^12–14^.

*Tet2* is abundantly expressed in the mouse brain ^10,15,16^ and through its DNA demethylation activity, it has been shown to regulate gene expression, neurogenesis, synaptic transmission, learning, and memory ^10,17^. In general, the TET enzymes influence cognitive function. While loss of *Tet1* interferes with memory extinction in young mice ^18^, loss of *Tet3* in hippocampal neurons increases excitatory glutamatergic synaptic transmission ^19^. Overexpression of neuronal *Tet2* also impairs hippocampal-dependent memory, while loss of *Tet2* in mature excitatory forebrain neurons enhances memory ^17^. It is unknown whether *TET2* regulates the expression of gene networks critical for PD progression or whether *TET2* controls pathways involved in the propagation of αSyn pathology.

Here, we examined gene networks and biological pathways altered in PD by performing unbiased RNA sequencing (RNAseq) analysis of neuronal nuclei isolated from the prefrontal cortex of PD patients relative to healthy controls. Cortical neurons show αSyn-rich Lewy pathology but do not degenerate early in PD cases and therefore remain present for investigation ^3^. We found a dysregulation of gene networks involved in known PD-associated pathways, such as oxidative phosphorylation, mitochondrial translation, autophagy, synaptic signaling, and intracellular transport. Intriguingly, we also found alterations in several genes associated with primary cilia. We then assessed the role of primary cilia in PD-relevant models and discovered that enhancing ciliogenesis ameliorates αSyn pathology *in vitro*. Additionally, we found that *Tet2* deletion also enhances ciliogenesis, while protecting against the spread of αSyn pathology and the degeneration of dopamine neurons *in vivo*.

## RESULTS

### Primary cilia gene networks are altered in PD neurons

To identify biological processes and gene networks altered in PD, we performed transcriptomic analysis. We compared RNAseq data from isolated neuronal nuclei in the prefrontal cortex of 105 individuals (26 early-stage PD patients, Braak stage 3-4; 27 late-stage PD patients, Braak stage 5-6; and 52 healthy controls) (Supplementary Table 1). A flow cytometry-based approach was used to isolate neuronal nuclei from the prefrontal cortex (Supplementary Fig. 1). After normalization and variance correction, we used a robust linear regression model to identify differentially expressed genes (DEGs) adjusting for age, sex, brain hemisphere, postmortem interval, sample origin, and possible sources of unwanted variation estimated with RUV. Relative to the matched healthy controls, we found 6,481 DEGs (*q* < 0.05) in the prefrontal cortex of patients with PD. Of these genes, 3,939 (60.8%) were downregulated while the remaining 2,542 (39.2%) were upregulated (Fig. 1a, b).

**Fig. 1.**
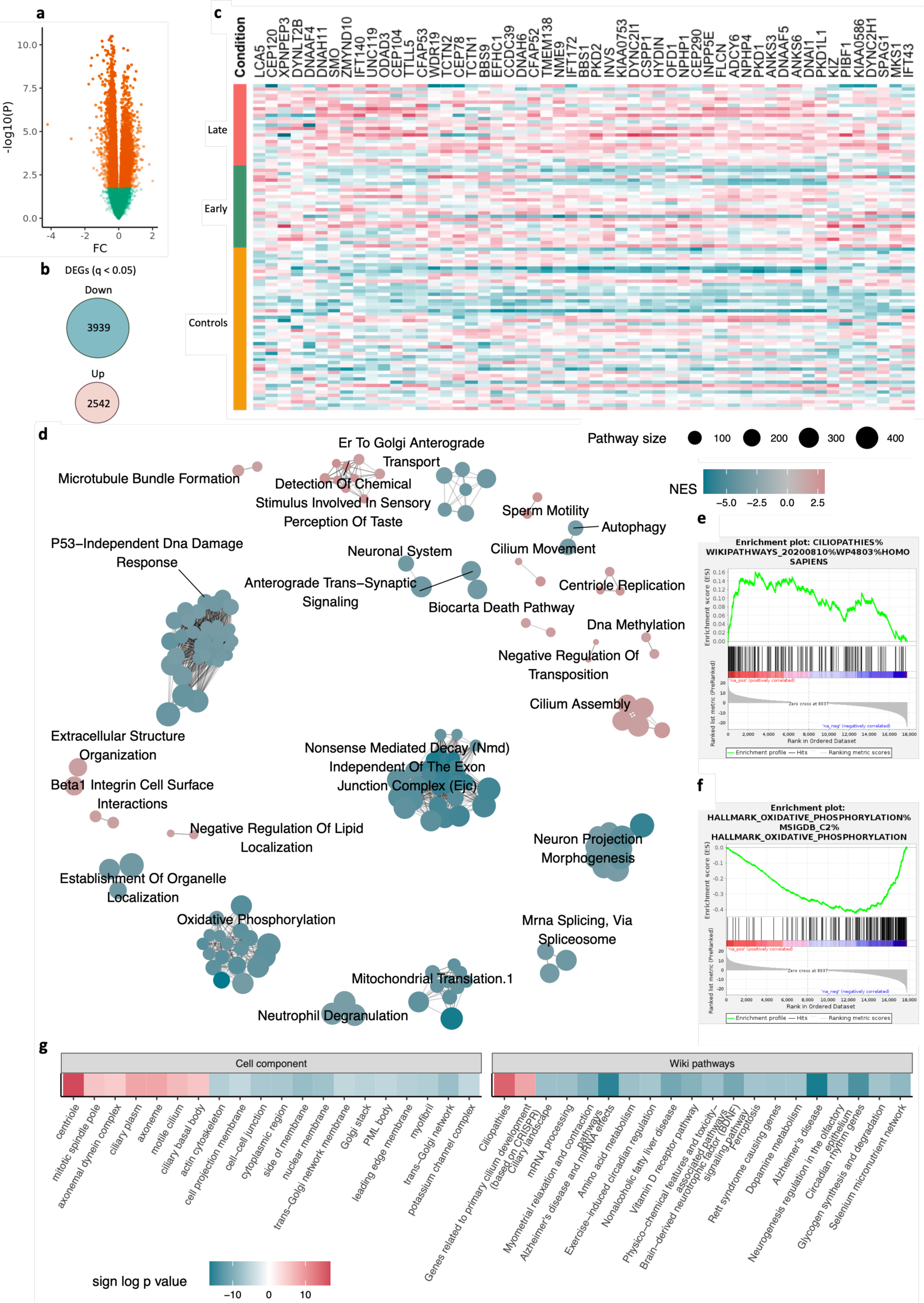
Dysregulation of primary cilia gene networks in cortical neurons from PD patients. Differentially expressed genes (DEGs) in the prefrontal cortex of PD neurons were identified in our RNAseq dataset using a robust linear regression model **a** Significantly altered genes in PD cortical neurons are highlighted in orange (*q* < 0.05, log(FC) ≥ 0.5, robust linear regression, multiple testing corrected). **b** Number of significantly dysregulated (up- and down-regulated) genes in all PD neurons relative to controls (*q* < 0.05). **c** Heatmap of dysregulated primary cilia-associated genes under the ‘ciliopathies’ hub. **d** Network and clustering of the significantly enriched pathways obtained via GSEA of the transcriptomic changes in PD neurons, with nodes representing altered pathways clustered into networks by EnrichmentMap and node colors constituting normalized enrichment score **e, f** Enrichment plots for ciliopathies (**e**) and oxidative phosphorylation (**f**), **g** Heatmap of dysregulated pathways in PD neurons as revealed by Cell component and Wiki pathways.

We then performed gene set enrichment analysis (GSEA) based on the identified DEGs using multiple pathway databases (including Cell component, Biological, and Wiki pathways) ^20,21^ and identified pathways with prominent gene network alterations in PD neuronal nuclei. Enriched pathways showed downregulation of oxidative phosphorylation, mitochondrial translation, autophagy, synaptic signaling and intracellular transport (Fig. 1d, f), supporting previous findings in PD ^22^. Intriguingly, we also identified alterations in primary cilia-associated gene networks, including upregulation of gene networks for cilium assembly (and ciliopathies), cilium movement, centriole replication and microtubule bundle formation, but a reduction of ciliary landscape in PD samples (Fig. 1d-g). Primary cilia are tiny hair-like microtubule-based structures that protrude from the surface of cells and function as signaling hubs ^23,24^. Further evaluation of the primary cilia alterations in the sorted PD neurons showed that the upregulation of primary cilia-associated gene networks began in early-stage patients with PD and persisted in late-stage patients (Supplementary Fig. 2).

To gain further insight into the extent of cilia-associated gene network alterations in PD neurons, we examined the genes altered under ‘Ciliopathies’ as defined in Wiki Pathways (Fig. 1c-e). A total of 53 upregulated genes formed the ‘Ciliopathies’ hub (Fig. 1c-e). Based on their localization and function, cilia associated genes can be categorized into specific groups: ciliogenesis (involved in cilia generation), axoneme (a microtubule-based cytoskeletal structure that forms the core of a cilium), intraflagellar transport (IFTs; involved in cargo transport along the axoneme), BBSome (a component of the basal body involved in cargo transport), RABs (RAB GTPases; involved in cargo transport and ciliary axoneme extension), and transition zone (associated with a specialized domain at the base of the cilium controlling protein entry and exit), as well as nexin-dyenin regulatory complex (involved in transport and motility of cilia) and SHH signaling (involved in regulating primary cilia signaling) (Fig. 1c) ^24–26^. Our analysis of the genes under ‘Ciliopathies’ revealed enhancement of genes associated with several of these categories. Specifically, we found increases in the expression of genes associated with ciliogenesis (including centriole or basal body genes), axoneme, transition zone, nexin-dynein complex, and SHH signaling (Fig. 1c).

Next, we generated heatmaps of DEGs commonly known to be associated with primary cilia based on their localization and function (Supplementary Fig. 3a-e). We observed an overrepresentation of upregulated genes belonging to ciliogenesis, axoneme, transition zone, and nexin-dynein regulatory complex (Supplementary Fig. 3a-e). Of particular interest was the upregulation of sonic hedgehog (SHH) signaling genes (*SMO, PTCH2, GLI4, KIAA0586*) in PD neurons relative to controls, as well as GLI-Similar 1 and 3 (*GLIS1* and *GLIS3)* genes (Supplementary Fig. 3a-e). *KIAA0586* (also known as *TALPID3*) is a crucial gene in primary cilia formation and hedgehog signaling, and its variation causes ciliopathies such as Joubert syndrome (an autosomal recessive neurological disorder) ^27,28^. SMO is also a key player in the SHH signaling pathway and its accumulation in primary cilia activates GLI transcription factors, which are the main mediators of primary cilia signal transduction ^29,30^. In contrast, genes associated with intracellular and ciliary trafficking (IFT, RAB and BBSome) were predominantly decreased (Supplementary Fig. 4), supporting previous findings in precursor PD neurons ^31^. Taken together, our transcriptomic analysis of cortical neuron nuclei sample sets revealed significant alterations in primary cilia gene networks in PD neurons from the frontal cortex relative to healthy controls, suggesting a potential role in PD.

### Ciliogenesis suppresses αSyn pathology *in vitro*

To further elucidate the involvement of primary cilia in PD, we utilized the well-established seed-based mouse primary cortical neuron model to evaluate how primary cilia enhancement affects αSyn pathology. Neurons were treated with αSyn preformed fibrils (PFF; 0.5 µg/mL) at 7 days *in vitro* (DIV) and fixed at 21 DIV. At the same time, neurons were treated with three different concentrations (0.5 μM, 1 μM and 3 μM) of purmophamine (PM), a compound known to activate ciliogenesis via the SHH signaling pathway ^32,33^. The test concentrations were not toxic to the neurons as purmophamine treatment did not significantly alter NeuN^+^ (neuronal nuclei marker) cell count or the area immunostaining positively for MAP2^+^ (neuronal dendrite marker) in comparison to results obtained from DMSO-treated controls (Fig. 2a-d). SHH signaling is initiated by the clearance of the receptor Ptch1, which leads to the ciliary accumulation of SMO and activation of Gli-dependent transcription ^29,30^, but purmophamine directly targets and activates SMO and primary cilia independent of Ptch1 ^32^. Here, two weeks of purmophamine treatment (at 0.5 μM, 1 μM and 3 μM doses) robustly increased the area covered by primary cilia in the primary cortical neurons, as indicated by the ACIII^+^ cilia (Fig. 2a, e). This increase in the area covered by primary cilia is likely due to the increased ciliary length observed in the purmophamine treated samples (Fig. 2a), rather than an increase in the number of primary cilia. To determine whether purmophamine treatment modulates αSyn pathology, we quantified the area covered by pS129 α-syn immunostaining in neurons treated with purmophamine compared to vehicle-treated controls. We observed dose-dependent reductions in αSyn pathology (7.6%, 14.9% and 23.8% in the 0.5 μM, 1 μM and 3 μM PM doses respectively), with the reduction at the 3 μM dose achieving statistical significance (*p* < 0.0001; one way ANOVA with Tukey’s multiple comparisons test; Fig. 2b, f). Together, these results suggest that boosting SHH-signaling enhances the area covered by primary cilia in cortical neurons, which attenuates the accumulation of pS129 αSyn pathology *in vitro*.

**Fig. 2.**
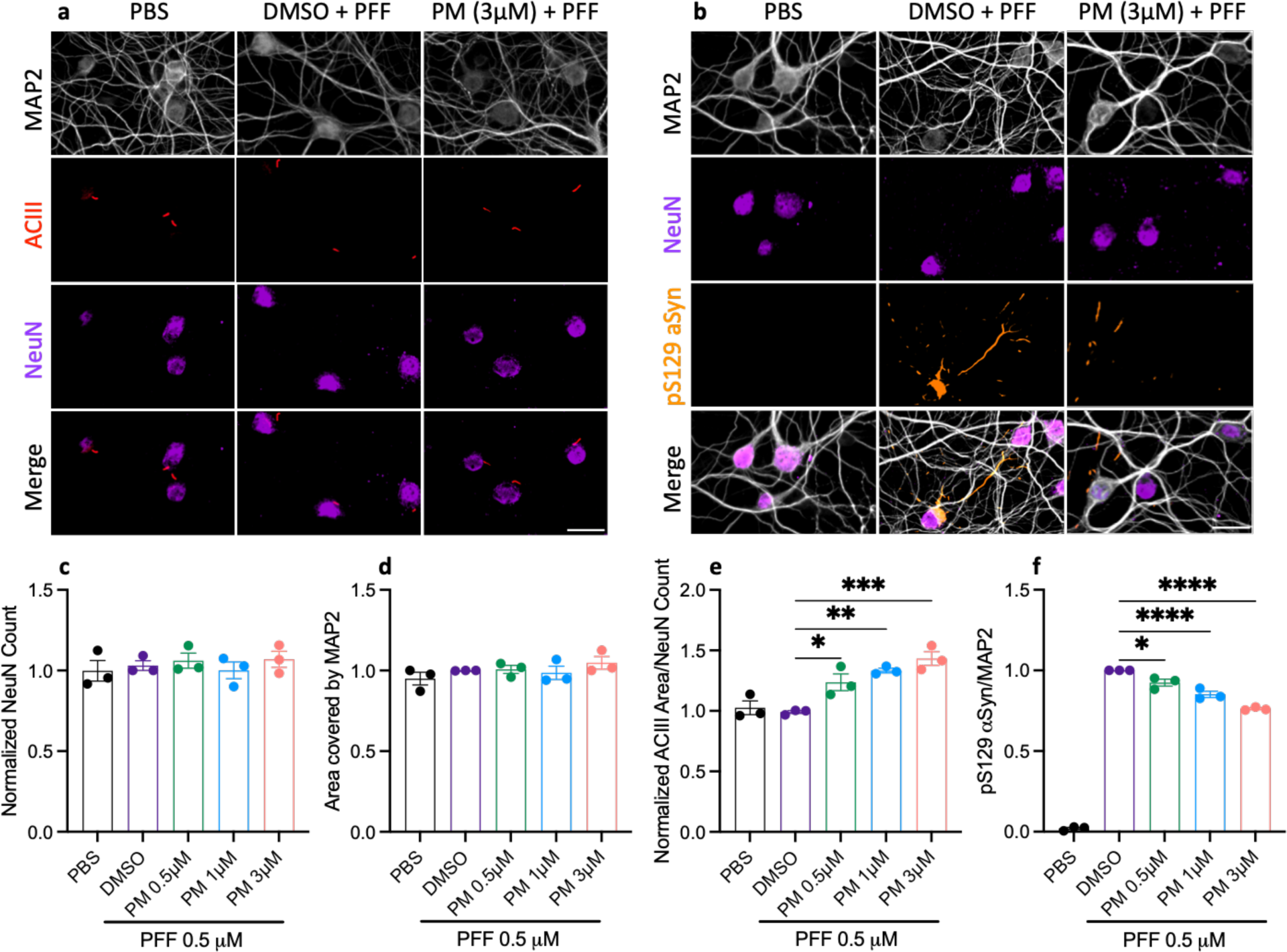
Enhancing ciliogenesis via SHH signaling suppresses αSyn pathology *in vitro*. Altered primary cilia and αSyn pathology in cortical neurons exposed to αSyn PFF for 14 days and incubated with different doses of purmophamine (PM). **a, b** Representative images showing ACIII^+^ (**a**) and pS129 αSyn (**b**) puncta (scale bar 50 µM). Analysis of **c** NeuN^+^ cortical neuron count normalized to vehicle (DMSO) control, **d** Area covered by MAP2^+^ neurons, **e** Area covered by ACIII^+^ primary cilia normalized to NeuN^+^ cell count, **f** pS129 αSyn pathology normalized to total MAP2 area. Data are mean ± SEM for 3 independent experiments combined, \**P* < 0.05, \*\**P* < 0.01, \*\*\**P* < 0.001, \*\*\*\**P* < 0.0001, one way ANOVA with Tukey’s multiple comparisons test relative to respective controls.

### *Tet2* inactivation suppresses αSyn pathology *in vivo*

Epigenetic mechanisms such as DNA methylation and chromatin/histone modifications are known to regulate pathways controlling ciliary biogenesis and cell cycle progression ^34^. TET2 is a master regulator of DNA methylation ^10^, and TET2-mediated DNA methylation abnormalities have previously been reported in PD ^8^. Interestingly, *Tet2* deletion appears to have beneficial effects in PD models ^8,15^, therefore we hypothesized that *Tet2* deletion might ameliorate αSyn pathology by stimulating genes associated with ciliogenesis. To test this hypothesis, we used a well-established mouse model of synucleinopathy in which intrastriatal injections of αSyn PFFs lead to widespread αSyn pathology ^35^. Using wildtype (WT) and *Tet2*^-/-^ mice, we examined whether knocking out *Tet2* affected the propagation of αSyn pathology by evaluating αSyn-immunostained sections 3- and 6-months after PFF injection. αSyn PFF injection in WT mice resulted in deposition of pS129 phosphorylated αSyn 3- and 6-months later (no αSyn pathology was detected in control mice injected with PBS; Fig. 3a-d). As expected, pS129-immunopositive αSyn pathology developed in the striatum and its anatomically interconnected regions like prefrontal cortex and substantia nigra at both timepoints (Fig. 3a-d). Interestingly, 3-months after αSyn PFF injection in *Tet2*^-/-^ mice, we found significantly less αSyn pathology in the cortex, striatum, and substantia nigra relative to WT mice (Fig. 3b). Indeed, there was 4- to 7-fold less αSyn pathology ipsilateral to the injection in each examined brain region when comparing to WT mice that received the same αSyn PFF dose (*p* < 0.0001, two-way ANOVA; Fig. 3b). Similar reductions in αSyn pathology were also observed contralateral to the injection in *Tet2*^-/-^ mice relative to WT mice 3-months post-injection (*p* < 0.0001, two-way ANOVA; Fig. 3b). Consistent with the data at the 3-month timepoint, we found significant reductions of αSyn pathology in all brain regions examined 6-months after PFF injection (*p* < 0.01, two-way ANOVA; Fig. 3c-d). In summary, *Tet2*^-/-^ mice exhibited lower seeding and propagation of αSyn pathology after injection of PFFs.

**Fig. 3.**
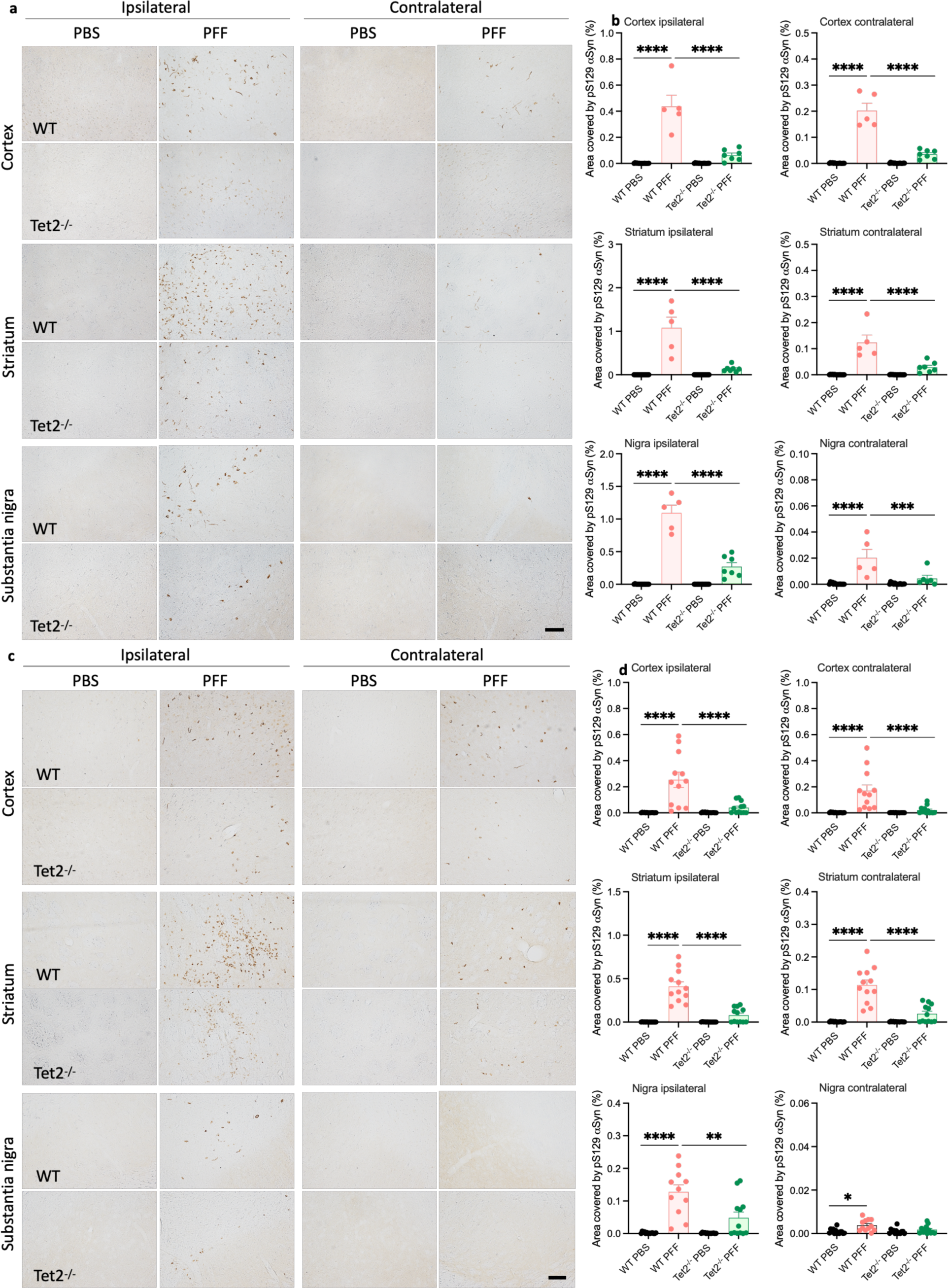
*Tet2* inactivation reduces αSyn pathology. Representative images (scale bar 250 µM) and quantification of pS129 αSyn pathology in the frontal cortex, striatum, and substantia nigra at **a, b** 3 months (*n*=5-7 mice per PFF group, *n*=15 mice per PBS group) and **c, d** 6 months (*n*=11-12 mice per PFF group, *n*=13-14 mice per PBS group) after αSyn PFF injection. The percentage area with pS129 αSyn immunoreactive staining was quantified using the Aiforia platform. Data are mean ± s.e.m., analyzed using two-way ANOVA with Tukey’s multiple comparisons test relative to the respective controls, \**P* < 0.05, \*\**P* < 0.01, \*\*\**P* < 0.001, \*\*\*\**P* < 0.0001.

### αSyn pathology regulates *Tet2* expression

Next, we examined whether αSyn pathology in mice influences the expression of the members of the TET family. Thus, we performed qPCR analysis of bulk cortical tissue isolated from the αSyn PFF-injected mice (at the 6-month timepoint) and compared them with PBS-injected control mice. To begin with, we confirmed the loss of *Tet2* mRNA in the knockout model, along with a mild reduction of *Tet3* but not *Tet1* (Fig. 4a-c). Interestingly, αSyn pathology reduced the mRNA expression of both *Tet2* and *Tet3* in WT mice, but not *Tet1* (Fig. 4a, b).

**Fig. 4.**
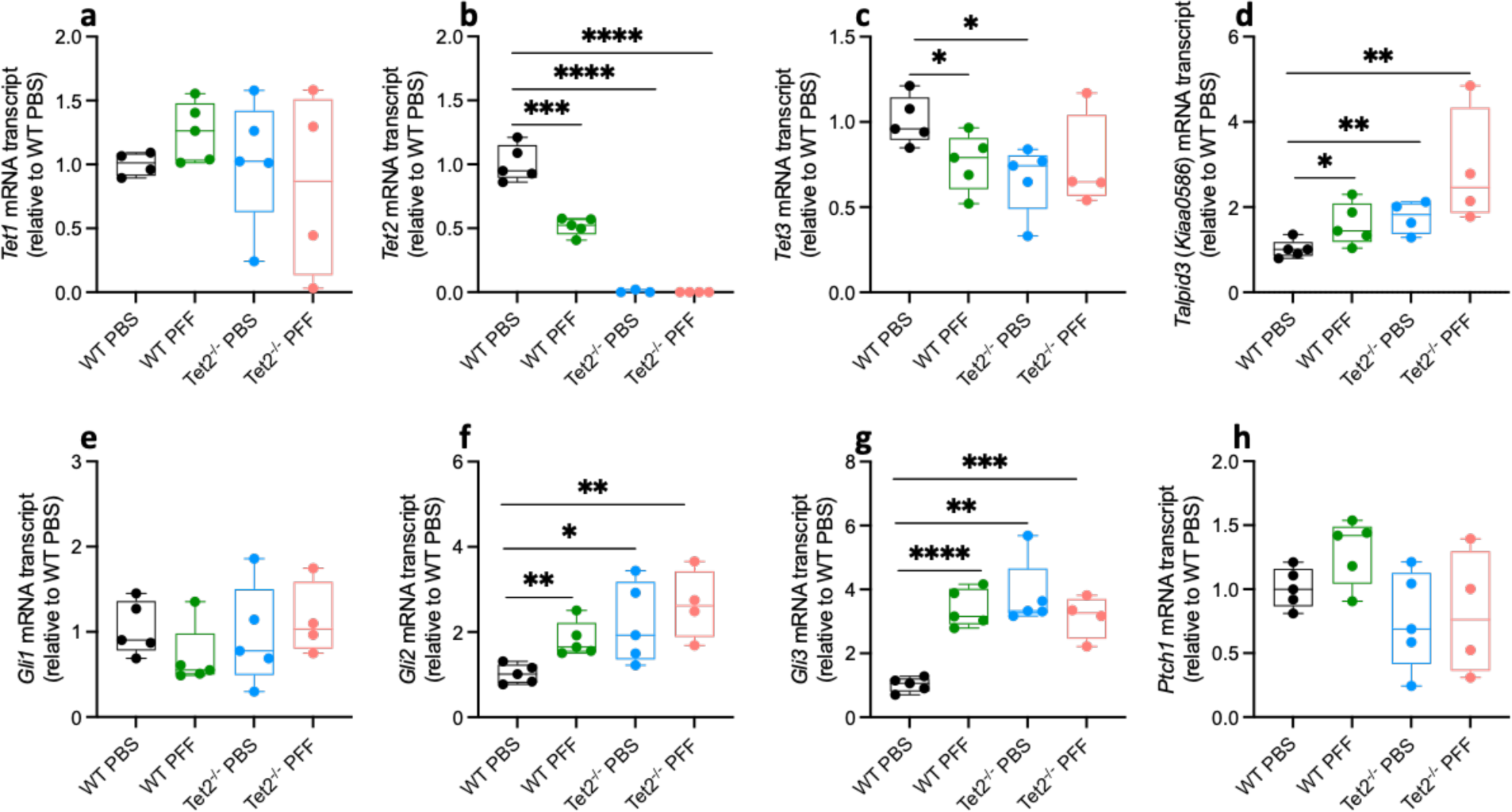
*Tet2* deletion enhances ciliogenesis via SHH signaling. Six months after striatal αSyn PFF inoculation, we examined the cortical expression of multiple genes using qPCR, **a** mRNA expression of *Tet1,* **b** *Tet2,* **c** *Tet3,* **d** *Talpid3,* **e***, Gli1,* **f** *Gli2,* **g** *Gli3,* and **h** *Ptch1* (*n*=3-5 mice per group). Data are mean ± s.e.m., analyzed using two-sided t-test, \**P* < 0.05, \*\**P* < 0.01, \*\*\**P* < 0.001, \*\*\*\**P* < 0.0001.

### *Tet2* inactivation and αSyn pathology stimulate ciliary gene expression

Next, we determined whether *Tet2*-knockout mediated suppression of αSyn pathology in our mouse model of synucleinopathy corresponds with changes in ciliogenesis. To evaluate the effect of *Tet2* inhibition on genes associated with primary cilia, we focused on the SHH signaling pathway. As already indicated, clearance of the SHH receptor Ptch1 causes the ciliary accumulation of SMO and the stimulation of Gli-dependent transcription ^29,30^. Gli1, Gli2 and Gli3 are the three main transcription factors that mediate SHH signaling. Gli1 is a transcriptional activator, but Gli2 and Gli3 can act either as activators or repressors ^30^. Gli2 and Gli3 are thought to be the most important regulators of SHH signaling target genes ^30^. Thus, we evaluated the mRNA expression of *Gli1, Gli2* and *Gli3,* as well as *Ptch1* in our *in vivo* model. Both *Tet2* deletion and αSyn pathology significantly enhanced the expression of *Gli2* and *Gli3* but not *Gli1* or *Ptch1* (Fig. 4e-h). *Talpid3* which had an increased expression in PD neurons (Fig. 1c; Supplementary Fig. 3; labelled as KIAA0586), was also significantly increased following *Tet2* deletion and αSyn PFF injection *in vivo* (Fig. 4d). Unlike their effect on these primary cilia and SHH-signaling genes, *Tet2* deletion and αSyn PFF did not significantly alter the mRNA expression of other genes such as *Tas2r13, Rnf139*, and *Phip* that are also dysregulated in PD neurons but are unrelated to primary cilia (Supplementary Fig. 5a-c). Together, these data show that the *Tet2*-knockout mediated suppression of αSyn pathology occurs together with an upregulation of ciliogenesis and SHH-signaling gene expression.

### *Tet2* deletion protects against αSyn-induced dopamine neuron loss

*Tet2* deletion has previously been shown to suppress inflammation and inflammation-induced nigral dopamine neuron degeneration in an LPS model of PD ^8^. Thus, we established whether *Tet2* deletion prevents αSyn pathology-induced inflammation and dopamine neuron degeneration. When we examined cytokine mRNA expression in our αSyn PFF mouse model at the 6-month timepoint, we found only modest evidence that αSyn pathology induced inflammation in the cortex. While the expression of *Il1b* was significantly increased in the cortex of WT mice, we did not detect changes in mRNA levels of *Il6, Il10,* or *Tnfa* (Supplementary Fig. 6a-d). Interestingly, *Tet2* deletion protected against the increased expression of *Il1b* (Supplementary Fig. 6a).

Given the accumulation and spread of αSyn pathology across multiple brain regions including the striatum and substantia nigra in our model, we assessed changes in dopamine neuronal count and any associated behavioral deficits. We evaluated tyrosine hydroxylase (TH)-immunostained sections in the substantia nigra 6 months after intrastriatal PFF injection. We found that along with the accumulation of αSyn pathology in the substantia nigra of WT mice 6 months after PFF injection, there was a 20% reduction in the number of TH-positive dopamine neurons (*p* = 0.0076, two-way ANOVA; Fig. 5a-d), without overt motor differences similar to previous reports ^35^. Importantly, we discovered that *Tet2* deletion mitigated loss of dopamine neurons (Fig. 5a, b). Together, our data shows that *Tet2* deletion is protective against αSyn-induced *Il1b* upregulation and dopamine neuron loss.

**Fig. 5.**
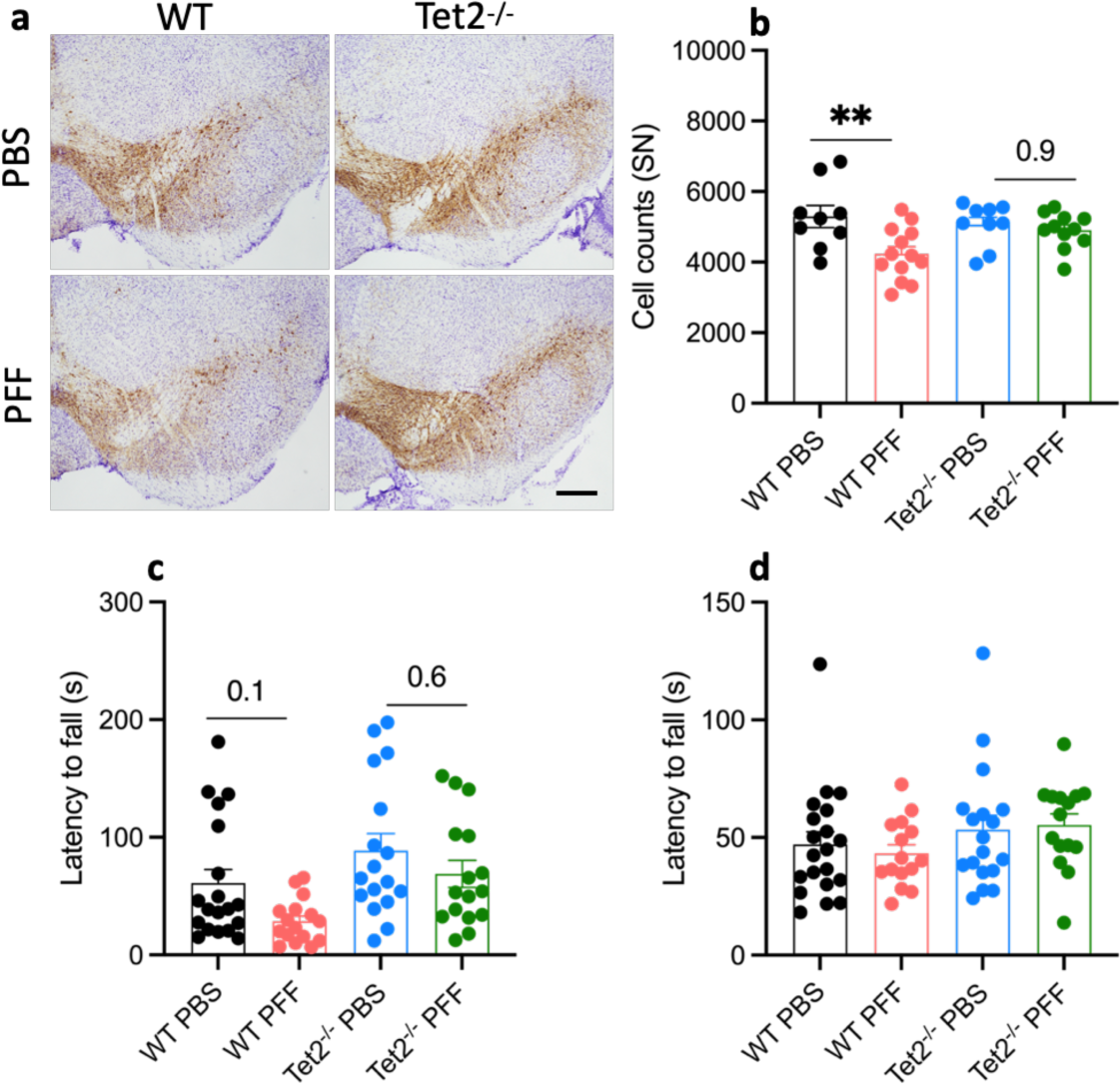
*Tet2* inactivation protects against αSyn-induced dopamine neuron loss. **a** Representative image (scale bar 500 μM) and **b** quantification of TH-positive neurons in the substantia nigra 6 months after αSyn PFF injection (*n*=11-13 mice per PFF group, *n*=9 mice per PBS group). TH immunoreactive staining was quantified using the Aiforia platform. **c, d** Behavioral assessment i.e., wire hang test (***c***) and rotarod (***d***) were performed 6 months after αSyn PFF injection (*n*=15-16 mice per PFF group, *n*=17-20 mice per PBS group). Data are mean ± s.e.m., analyzed using two-way ANOVA with Tukey’s multiple comparisons test relative to the respective controls, \*\**P* < 0.01.

## DISCUSSION

We performed RNAseq analysis of neuronal nuclei isolated from postmortem cortical samples in early and late-stage PD patients and found 6,481 dysregulated genes (60.8% of them downregulated) in PD samples relative to controls. Previous studies performing bulk RNAseq analysis of cortical, striatal or nigral tissues from PD patients have reported similar percentages of downregulated genes in PD ^31,36,37^. Gene expression profiles in bulk brain tissues are influenced dramatically by differences in cellular composition ^38^. Because our method is focused on neuronal nuclei, it provides more insight into changes occurring specifically in neurons. Interestingly, we found that major PD-associated phenotypes such as oxidative phosphorylation, mitochondrial translation, autophagy, synaptic signaling, and intracellular transport are all impaired in the prefrontal cortex of PD patients, processes which have previously been considered to be impacted by PD pathogenesis ^37,39^.

A major finding of our unbiased molecular characterization of postmortem cortical neuronal nuclei was the alteration of genes associated with primary cilia in both early and late-stage PD brains. Specifically, we found significant upregulation of primary cilia gene networks controlling ciliogenesis, axoneme, transition zone shuttling, SHH signaling, and nexin-dynein regulatory complex, along with a downregulation of intraflagellar or ciliary transport (Fig. 1; Supplementary Fig. 3, 4). The downregulation of intraflagellar and ciliary transport genes is suggestive of defective trafficking and appears to be consistent with a prior study that reported reductions in genes controlling ciliary transport in the substantia nigra of PD cases ^31^. However, in contrast to our findings, that study also reported reductions in genes regulating primary cilia length and transition zone shuttling in bulk substantia nigra tissues from late-stage sporadic PD patients ^31^. Given that we conducted RNAseq on sorted cortical neurons from early and late-stage patients with PD instead of bulk nigral tissues, our data likely reflect changes occurring specifically in cortical neurons in PD patients. Alterations in primary cilia-associated gene networks are known to induce changes in ciliary density, morphology and function ^24^. Interestingly, the upregulated ciliary gene networks observed in this study are consistent with previous reports showing enhanced ciliogenesis including increased primary cilia density and length in sporadic PD patients and in mouse (i.e., 6-OHDA and MPTP) models of PD ^31,40,41^.

Projecting on neurons in the mature brain, primary cilia act as antennas that detect extracellular (including hedgehog) signals, with additional roles in transducing detected signals to regulate cellular processes (such as excitatory synapses and neurogenesis) ^42,43^. The reaction of primary cilia to changes in their environment happens in a time and tissue-dependent manner ^24,26^. As such, the observed changes in primary cilia-associated gene expression and ciliary morphology could either be driven directly by PD pathogenic processes or they could represent compensatory mechanisms that occur in response to the core pathogenesis. In PD, disease-associated motor symptoms (leading to clinical diagnosis) are believed not to occur until Braak stages 3-4 when αSyn-rich Lewy pathology reaches the substantia nigra and there is significant (typically estimated at 50-70%) loss of dopaminergic neurons. At Braak stages 3-4, Lewy pathology has not yet affected the neocortex (this occurs in the cortex at Braak stages 5-6 i.e., late-stage PD) ^3^. Since the observed primary cilia gene network alterations in cortical neurons were both in the early (Braak stages 3-4) and late-stage (Braak stages 5-6) PD patient samples, we propose that these ciliary gene network alterations are a compensatory reaction to *prolonged* exposure to the molecular events that are integral to PD pathogenesis. We tested this hypothesis by exposing primary cortical neurons to αSyn PFF acutely. In line with the notion that prolonged exposure to PD pathogenesis is necessary, we found no changes in ciliation in this paradigm (Fig. 2). In contrast, when we examined neurons in WT mice that had been subjected to prolonged (i.e., 6 months) exposure to αSyn PFFs, we did observe increased primary cilia and SHH signaling genes (Fig. 4), akin to the observation in cortical neurons of PD patients (Fig. 1; Supplementary Fig. 3). Interestingly, at the 6-month timepoint, the observed αSyn pathology in the WT mice was also associated with significant loss of nigral dopamine neurons (Fig. 5). Loss of midbrain dopaminergic input has previously been reported to alter ciliary dynamics by increasing primary cilia length ^40,41^.

To examine the role of a compensatory enhancement of ciliogenesis in a cell culture model of PD involving exposure to αSyn PFFs, we induced ciliogenesis in primary cortical neurons with the SHH-signaling activator purmophamine. Interestingly, we found that ciliogenesis significantly suppressed the accumulation of αSyn pathology in the cortical neurons (Fig. 2). In analogy to the present finding, previous studies have shown that enhancing ciliogenesis (e.g., via SHH signaling) protects adult dopamine neurons against acute neurotoxic insults (i.e., MPTP, 6-OHDA and rotenone)^41,44,45^. The mechanism whereby ciliogenesis suppresses the accumulation of αSyn pathology and protects adult dopamine neurons against acute neurotoxic insults remains unclear. Importantly, enhancing ciliogenesis (e.g., through purmophamine treatment, *ptch1* knockout, or overexpression of GLI’s) increases autophagy ^41,46^, which might be one explanation why ciliogenesis suppresses the accumulation of αSyn pathology.

DNA methylation regulates ciliary biogenesis, together with other processes such as cell cycle progression ^34^. Therefore, we also investigated the effect of the epigenetic regulator *Tet2* on ciliogenesis. We found that mice lacking *Tet2,* relative to WT mice, exhibited increased expression of primary cilia- and SHH-signaling genes (i.e., *Gli2*, *Gli3* and *Talpid3)*. This increase in primary cilia- and SHH-signaling genes was more pronounced in *Tet2* knockout mice analyzed 6 months after exposure to αSyn PFF (compared to similar mice not injected with αSyn PFFs). Interestingly, in addition to enhancing cilia-associated genes, *Tet2* deletion was also associated with reduced spread of αSyn pathology and significantly less dopamine neuron degeneration, in analogy to prior reports of reduced nigral neuron death in MPTP and LPS models of PD applied to *Tet2* knockout mice ^8,9^. The enhanced expression of cilia-associated genes in *Tet2* null mice and the reduced nigral neurodegeneration in αSyn, LPS and MPTP models of PD, suggests that increased ciliogenesis might contribute to the protective effect of *Tet2* deletion.

In summary, our study uncovers a convergence between the epigenetic marker *Tet2* and ciliogenesis in the modulation of αSyn pathology. We discovered that PD cortical neurons are associated with profound changes in primary cilia gene networks. Enhancing ciliogenesis via SHH signaling in primary cortical neurons increased neuronal ciliation, which suppressed the accumulation of αSyn pathology. Interestingly, *Tet2* deletion also enhanced primary cilia gene networks and rescued the accumulation of αSyn pathology and dopamine neuron degeneration. Future studies must investigate brain region and cell-type specific adaptations in ciliary gene expression in PD and should explore the mechanisms underlying ciliogenesis-induced suppression of αSyn pathology.

## METHODS

### Human cortical samples

All experiments involving the use of postmortem brain samples were done with approval from the ethics committee of the Van Andel Institute (Institutional Review Board number 15025). The prefrontal cortex samples were obtained with written permission from the Michigan Brain Bank, the Parkinson’s UK Brain Bank, and the NIH NeuroBioBank. For each of the brain samples used in this study, we have provided information on clinical variables (PD diagnosis and Braak stage), sample origin, postmortem interval, brain hemisphere, and patient demographics (age and gender). The full patient data can be found in Supplementary Table 1. Overall, cortical samples used in this study were from PD patients with confirmed brain Lewy pathology, while controls had no Lewy pathology and were pathologically normal. Neurons from the prefrontal cortex were used in this study since: (a) this brain region plays a critical role in the pathophysiology of PD, with Lewy pathology spreading to this region ^3^, and (b) neurons remain present in the cortex unlike those in the substantia nigra which undergo severe degeneration ^47^. We isolated neuronal nuclei from the prefrontal cortex samples using a flow cytometry-based approach, as previously described ^8,48,49^. A description of the gating strategy used has been provided in Supplementary Fig. 1. Using this gating strategy, we sorted approximately 200,000 NeuN^+^ nuclei for each cortical sample, and then performed RNAseq analysis.

### RNA sequencing of human cortical samples

After sorting the NeuN^+^ nuclei, RNA was isolated from neuronal nuclei using the RNeasy Micro Kit (Qiagen, 74004). The quantity and quality of RNA were assessed by Qubit RNA High Sensitivity Kit (Thermo Fisher Scientific, Q32852) and Bioanalyzer (Agilent Technologies). RNA library preparation and sequencing were performed at the Van Andel Institute Genomics Core. Libraries were generated using the SMarter® Stranded Total RNA-Seq Kit v3 – Pico Input Mammalian (Takara, 634488) and sequenced on a NovaSeq6000 Sequencer with 2x100 bp reads and a minimum of 50 million reads per sample. Base calling was done using Illumina RTA3 and the output of NCS demultiplexed and converted to FastQ format with Illumina Bcl2fastq (v1.9.0).

We used TrimGalore (v.0.4.4) to remove adapter sequences in our fastq data files. We then employed STAR (v.2.3.5a) to align and count sequencing reads to the human (GRCh37/hg19) Gencode v19 primary assembly transcriptome. TMM normalization was performed in edgeR (v.3.24.3) ^50^. Prior to this normalization, the gene counts matrix was imported into R (3.5.1) and the genes with low expression (<1 million in all samples) were removed. CIBERSORT ^51^ (https://cibersortx.stanford.edu/; 100 permutations) was used to perform cell-type deconvolution with 834 brain cell-type-specific gene signatures ^52^. To identify differentially expressed genes in PD relative to controls, we used robust linear regression or negative binomial models implemented in limma (v.3.50.1) after variance adjustment with the help of voom package. We adjusted for age, brain hemisphere, sex, sample origin and sources of unknown variation. We determined the sources of unknown variation using stable control genes (*p* ≥ 0.5) in the RUVseq Bioconductor package (v.1.28.0) ^53^. Benjamini-Hochberg correction for multiple testing and log(FC) threshold (with significance set at *q* < 0.05 and absolute log(FC) 2 0.5) were applied.

Next, we performed pathway enrichment analysis using the GSEA (v4.1.0) software (https://www.gsea-msigdb.org/gsea/index.jsp). We ranked genes according to their fold changes (whether upregulated or downregulated), and their negative log transformed *p* values. Next, we obtained the human gene set files from Bader lab website ^54^ in the last quarter of 2022 and performed 1,000 permutations to obtain pathway enrichment *p* values. The most significantly dysregulated (top 50 up-regulated and top 50 down-regulated) pathways were clustered together and their interaction networks were visualized using the aPEAR package ^55^. Naming of the clusters was accomplished using the most influential pathway within each cluster, as determined by PageRank algorithm. We also used clusterProfiler in R package to obtain the word cloud of the most significant pathways.

### Generation of αSyn PFF

αSyn was expressed from plasmid (Addgene, 89075) in BL21 (DE3) bacterial strain after ampicillin selection. In this study, we used pD454-SR mouse alpha-synuclein (Addgene plasmid, 89075; http://n2t.net/addgene:89075; RRID: Addgene_89075). Large 250 ml bacterial cultures were left overnight at 37°C with constant shaking at 600 rpm to allow for αSyn expression.

Bacteria were harvested after reaching 0.6-0.8 OD 600, the pellets were resuspended in high salt buffer (750 mM NaCl, 10 mM Tris at pH 7.6, 1 mM EDTA and 1 mM PMSF), and homogenized by sonication for a total of 80 s (5 s on and 25 s off). The homogenate was briefly boiled, and the lysate cleared by centrifugation at 10,000 g. Monomeric αSyn was purified by size exclusion chromatography using a Superdex 200 column (Cytiva) previously equilibrated in size exclusion buffer (50 mM NaCl, 10 mM Tris at pH 7.5 and 1 mM EDTA). Fractions were collected and analyzed by SDS-PAGE and Coomassie blue staining to ensure high levels of expression and exclusion of truncated forms of αSyn. Fractions corresponding to the desired αSyn were dialyzed O/N in 4 l of size exclusion buffer. Following dialysis, the fractions were concentrated using 5 kDa MWCO Amicon. Next, we performed anion exchange using a monotrap Q 5 ml column (Cytiva) and collected the desired αSyn fraction for SDS-PAGE and Coomassie blue staining. Endotoxin load was determined using GenScript Toxin Sensor Chromogenic LAL endotoxin assay kit following manufacturer’s instructions (L00350). The protein concentration was determined using BCA protein assay kit (Pierce, A53226) according to manufacturer’s instructions. PFFs were generated by incubation at 37°C for 7 days with constant shaking at 1,000 rpm (Eppendorf Thermomixer). After ultracentrifugation for 1 h at 100,000 g, the relative amount of PFFs produced was determined by comparing the amount of αSyn in the pellet to the soluble fraction.

### Stereotactic injections of αSyn PFF

Male and female C57BL/6J (000664) and Tet2^+/-^ mice (023359) obtained from The Jackson Laboratory were bred at Van Andel Institute’s vivarium. The mice were housed up to 4 mice per cage in a 12 h light/dark cycle with free access to food and water. All animal experiments were performed in accordance with the Guide for the Care and Use of Laboratory Animals (National Institutes of Health, United States) and were approved by the Animal Care and Use Committee at Van Andel Institute. To induce PD-like pathology, 8-12 weeks old C57BL/6J and *Tet2^-/-^* mice were anaesthetized with isoflurane/oxygen, placed in a stereotactic head frame (Stoelting, IL, USA), and injected with a local anesthetic (0.03 ml of 0.5% Ropivacaine; Henry Schein) at the site of incision. Injections were done using a 30-gauge needle and a glass capillary attached to a 10 μl Hamilton syringe. 2 μl of PBS or αSyn PFF (5 μg/μl in sterile PBS) were unilaterally injected in the striatum (coordinates: AP, +0.2 mm; L, ±2.0 mm; DV, -2.6 mm relative to bregma and dura) at a flow rate of 0.4 μl per minute. Following injection, the capillary was left in place for 3 min before being slowly removed. The mice were then given 0.5 ml saline and 0.04 ml Buprenex for hydration and pain relief, respectively. We returned the mice to their home cages until tissue collection.

### Behavioral testing

#### Rotarod

To assess motor coordination, we used an accelerating rotarod device (Ugo Basile, Italy). The mice had two days of training prior to the test sessions. During the training sessions, the mice were placed on a still rod, with the rod subsequently accelerated to 4 rpm on day 1 and 16 rpm on day 2. On day 3, the mice had three test sessions, with each session lasting up to 10 min. The mice were allowed to rest at least one hour between each test session. For the test sessions, the rod was accelerated from 4 to 40 rpms and the latency to fall was recorded (average of the three tests). The time was also stopped if a mouse gripped the rod and rotated with it rather than walking on it.

#### Wire hang test

Wire hang test (San Diego Instruments) was used to assess muscle strength and coordination. The mice had two test sessions, one per day. The fore limbs of the mice were placed on a suspended wire (35 cm high) over a soft-landing area for a maximum of 5 min. The latency to fall from the suspended wire was recorded (average of the two tests). The time was also stopped if a mouse balanced on or walked on the wire instead of hanging.

### Histology and image analysis

Three and six months after αSyn PFF injection, the mice were transcardially perfused using 0.9% saline followed by 4% paraformaldehyde (PFA) in phosphate buffer. The brains were post-fixed for 24 h in 4% PFA, switched to 30% sucrose in phosphate buffer and kept at 4°C until sectioning. During sectioning, a sliding microtome was used to cut the brains into 40-μm free-floating coronal slices spaced by 240 μm for analysis. The sectioned tissues were incubated with anti-pSer129 αSyn (Abcam, Ab51253) or anti-TH (Millipore Sigma, 657012) sera overnight at room temperature, followed by a 2 h incubation with goat anti-rabbit secondary biotinylated antibody. To detect the antibody staining, we employed a standard peroxidase-based method (Vectastain ABC kit; Vector Laboratories), followed by incubation with 3, 3’-diaminobenzidine (DAB kit; Vector Laboratories). After placing the immunostained tissues on glass slides, they were dehydrated and coverslipped using the Cytoseal 60 mounting medium (Thermo Fisher Scientific). Using a ZEISS Axio Scan Z1 Digital Slide Scanner (Carl Zeiss Microscopy), the slides were scanned to generate a z-stack with a 3 μm scanning interval. For the blind-coded scanned images of the antibody-stained coronal sections, we analyzed pSer129 αSyn pathology, or TH^+^ dopamine neuron density using deep learning convolution neural networks and a supervised learning model developed by Aiforia Technologies ^56^.

### mRNA expression

Total RNA was isolated from mouse cortical samples using RNeasy plus mini kit (Qiagen). SuperScript III reverse transcriptase (ThermoFisher) was used to synthesize cDNA from the total RNA. Quantitative real-time PCR was then performed for the cDNA samples using SYBR Green Master mix (Roche) on an Applied Biosystems StepOnePlus real-time PCR machine (Thermo Fisher). ΔΔCT values for target genes were relative to *Gapdh*. Primers used in detecting the expression levels of the target genes are as listed in Supplementary Table 2.

### Primary neuron culture

Primary neuron cultures were prepared from embryonic day 18 (E18) CD1 mice (strain 022; RRID: IMSR_CRL:022). Brains were gently removed from the embryos and placed into a petri dish filled with ice-cold, sterile Hibernate Medium (Gibco, A1247601). Brain hemispheres were gently separated, and cortical tissue isolated from each hemisphere and pooled. Pooled cortical tissues were digested in papain solution (Worthington, LS003126, 20 U/mL) and then treated with DNase I (Worthington, LS006563) to remove residual DNA. The tissue was then washed with pre-warmed Neurobasal media (Gibco, 21103049), mechanically dissociated, and strained through a 40 μm cell strainer. The cell suspension was pelleted at 1000 rpm, resuspended in Neurobasal media (containing 1% B27, 2 mM GlutaMAX, and penicillin-streptomycin), and gently mixed. The dissociated neurons were seeded on poly-D-lysine (Sigma, P0899) coated 96 well culture plates (Perkin Elmer, 6005182) at 17000 cells/well and maintained at 37°C at 5% CO2. The cells were treated with α-syn PFF (0.5 μg/ml) and different concentrations (i.e., 0.5 μM, 1 μM and 3 μM) of purmophamine along with its respective controls at DIV 7. The cells were maintained till DIV 21 followed by immunocytochemistry. Three independent experiments were performed with three technical replicates in each experiment.

### Immunofluorescence staining

Cortical neurons were initially fixed with prewarmed 4% PFA/4% sucrose in phosphate buffered saline (PBS) and then washed with PBS. The cells were then permeabilized with 0.03% Triton X-100 in 3% bovine serum albumin (BSA) and washed with PBS. Cells were blocked with 3% BSA for 1 hour before overnight incubation with primary antibodies at 4°C. The following primary antibodies were used: pS129 α-synuclein (Biolegend, 825701; 1:2000), MAP2 (Abcam, AB5543; 1:5000), NeuN (Millipore, MAB377; 1:1500), ACIII (EnCor, RPCA-ACIII 1:1000). Cells were then incubated with respective AlexaFluor (Invitrogen) secondary antibodies for 1 hour at room temperature with shaking, and then washed with PBS. The cells were then incubated with DAPI 1:10,000 in PBS, and plates were sealed and kept at 4°C for imaging. The plates were imaged at 20x zoom with a 0.5 optical for ten ROIs per well with Zeiss Cell discoverer 7 and analyzed in Zen 3.0 software. Area of pS129 α-synuclein, MAP2, ACIII along with counts of NeuN positive cells were quantified in a blinded manner for each ROI and pooled for each experimental group. pS129 α-synuclein area was normalized to total MAP2 area to determine pathology and the area covered by ACIII-positive signals was normalized to NeuN counts to determine the total area covered by ACIII-positive primary cilia for each experimental group. Data was normalized to the respective control group.

### Statistical analysis

Differentially expressed genes from our RNAseq dataset were analyzed using robust linear regression or negative binomial regression implemented in limma (v.3.50.1), followed by Benjamini-Hochberg correction for multiple testing (significance threshold set at *q* < 0.05). We applied Fisher’s exact tests for enrichment analyses. GSEA was used for pathway enrichment analysis and was corrected for multiple testing with the Benjamini-Hochberg method. Primary cortical neuron fluorescence data were analyzed using one way ANOVA with Tukey’s multiple comparisons test. Given the sample sizes, qPCR data comparing gene expression were analyzed using a two-sided t-test. Immunohistochemistry and behavioral studies in mice were analyzed using a two-way ANOVA with genotype and treatment as between-subjects factors, followed by Tukey’s multiple comparisons test. Data are presented as mean ± s.e.m.

## Data availability

All data generated or analyzed for this study are included in this published article and its Supplementary Information.

## Conflicts of interest

PB declares employment for F Hoffmann-La Roche and stock ownership for F Hoffmann-La Roche, Acousort AB, Axial Therapeutics, Enterin Inc and Kenai Therapeutics.

## Acknowledgements

This study was supported by a grant from Michael J. Fox Foundation under award number MJFF-001209 to P. B; transferred to L.B. subsequent to P.B.’s departure from VAI. We thank the VAI Vivarium, Genomics, and Optical Imaging Cores for technical assistance.

## Contributions

E.Q., M.L.E.G., M.H., P.B., L.B., L.L.M, and J.G., conceptualized, designed, and planned the experiments. E.Q., N.V., E.E., J.B., T.C., E.K., A.L., and C.G., performed the experiments. M.A., and E.Q., produced and validated the α-synuclein preformed fibrils. J.A.S., performed project administration. M.M., L.L.M., and J.G., performed statistical analyses.

## Ethics declaration

### Competing interests

The authors declare no competing interests.

---

**Supplementary Fig. 1.**
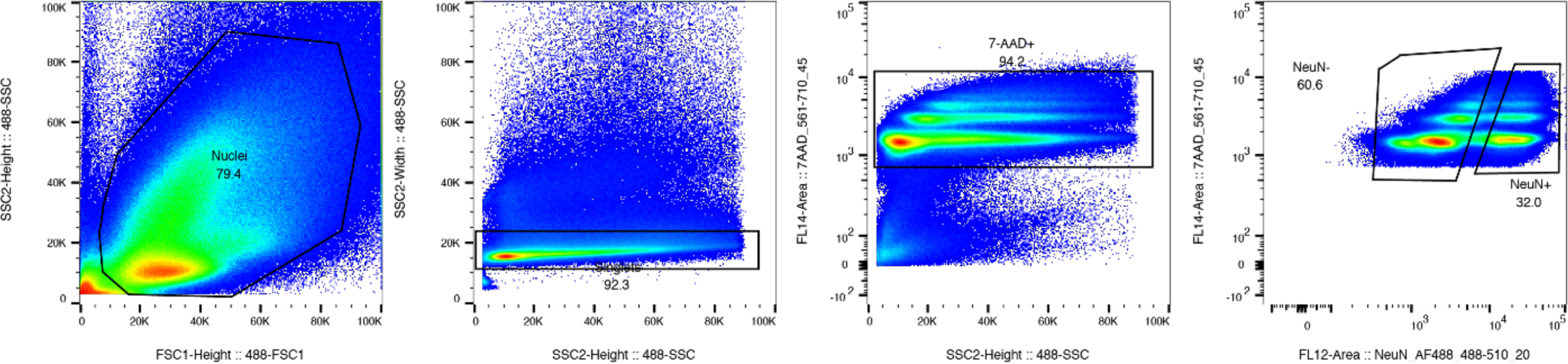
Flow cytometry and gating strategy. Gating strategy to isolate NeuN^+^ (NeuN and 7AAD stained) neurons from the prefrontal cortex of postmortem PD patient and control samples.

**Supplementary Fig. 2.**
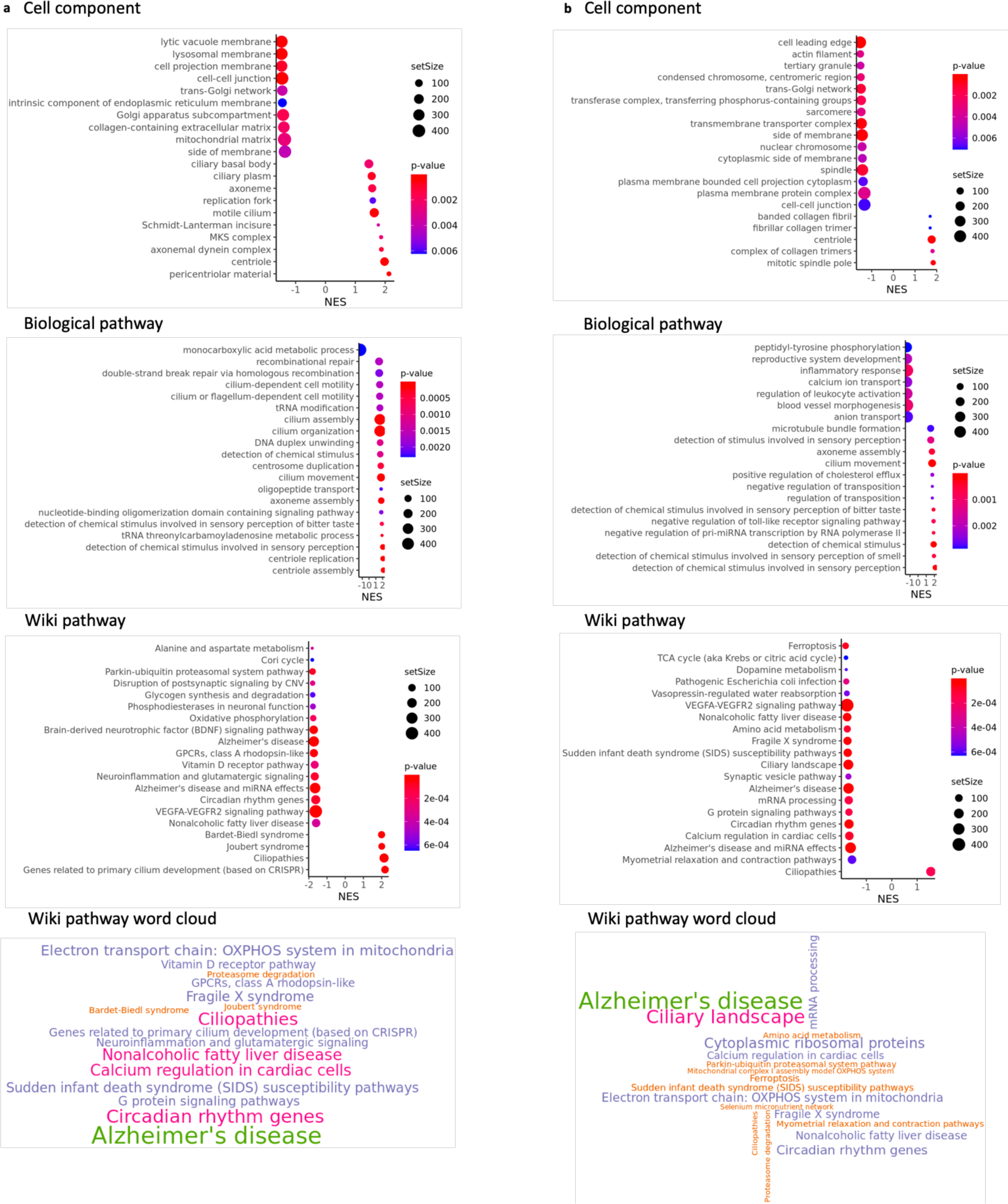
Top 20 dysregulated pathways associated with early and late-stage PD. Dysregulated pathways in PD prefrontal cortex neurons were identified in our RNA-sequencing dataset using a robust linear regression model. **a, b** Pathways affected by the transcriptomic changes in cortical neurons in the Cell component, Biological and Wiki pathways in early (**a**) and late-stage (**b**) PD patients, together with word cloud of significantly enriched Wiki pathways obtained through GSEA (i.e., gene set enrichment analysis).

**Supplementary Fig. 3.**
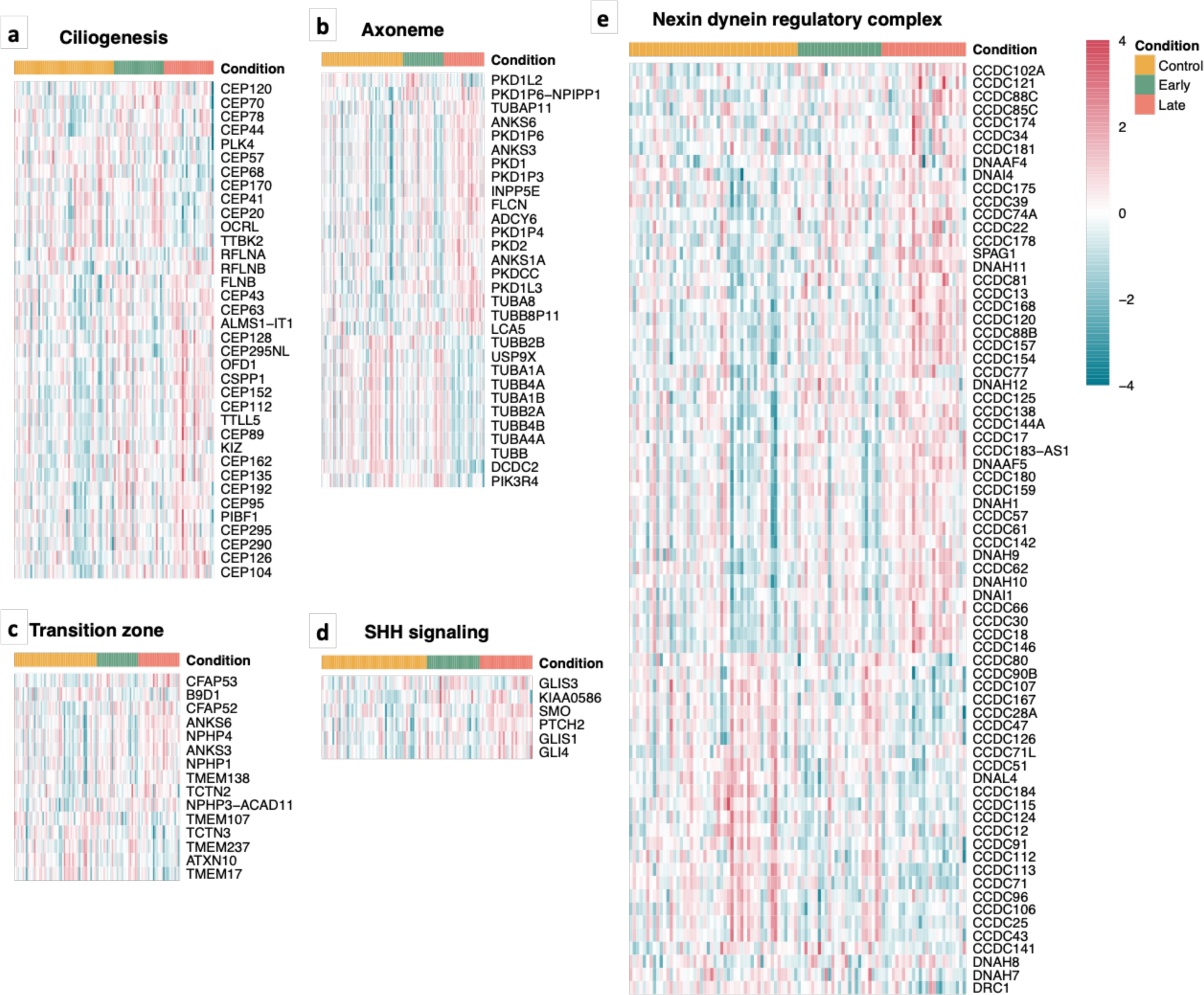
Enhanced ciliogenesis and SHH signaling genes in cortical neurons from PD patients. Altered gene networks in cortical neurons from PD patients were identified in our RNA-sequencing dataset using a robust linear regression model. **a-e** Heatmaps of primary cilia-associated genes belonging to ciliogenesis (**a**), axoneme (**b**), transition zone (**c**), SHH signaling (**d**), and nexin-dynein regulatory complex (**e**).

**Supplementary Fig. 4.**
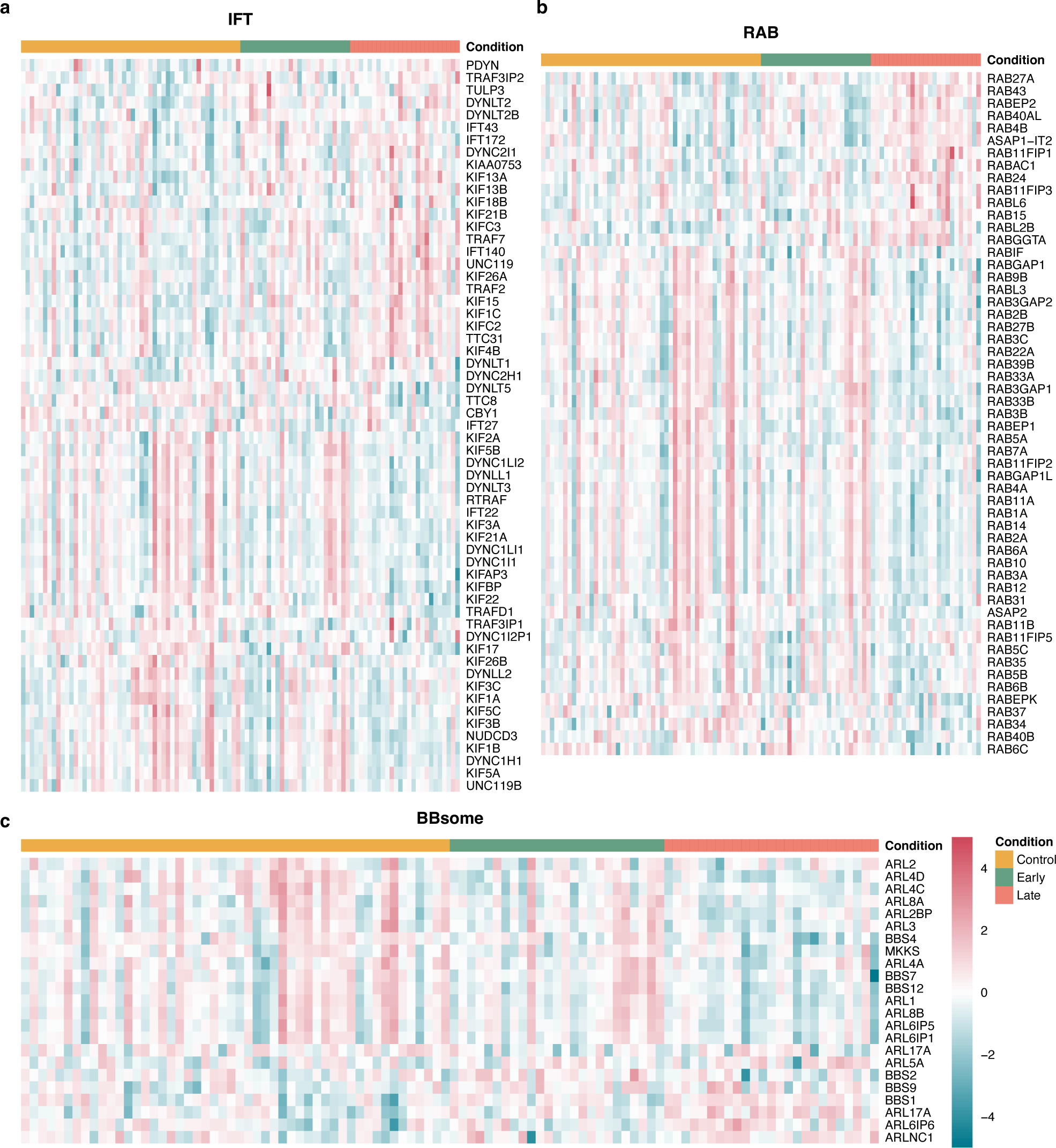
Downregulation of intraflagellar, BBSome and RAB genes in cortical neurons from PD patients. Dysregulated pathways in PD cortical neurons were identified in our RNA-sequencing dataset using a robust linear regression model. **a-c** Heatmaps of dysregulated primary cilia-associated genes belonging to intraflagellar (IFT) (**a**), BBSome (**b**), and RAB (**c**) categories.

**Supplementary Fig. 5.**
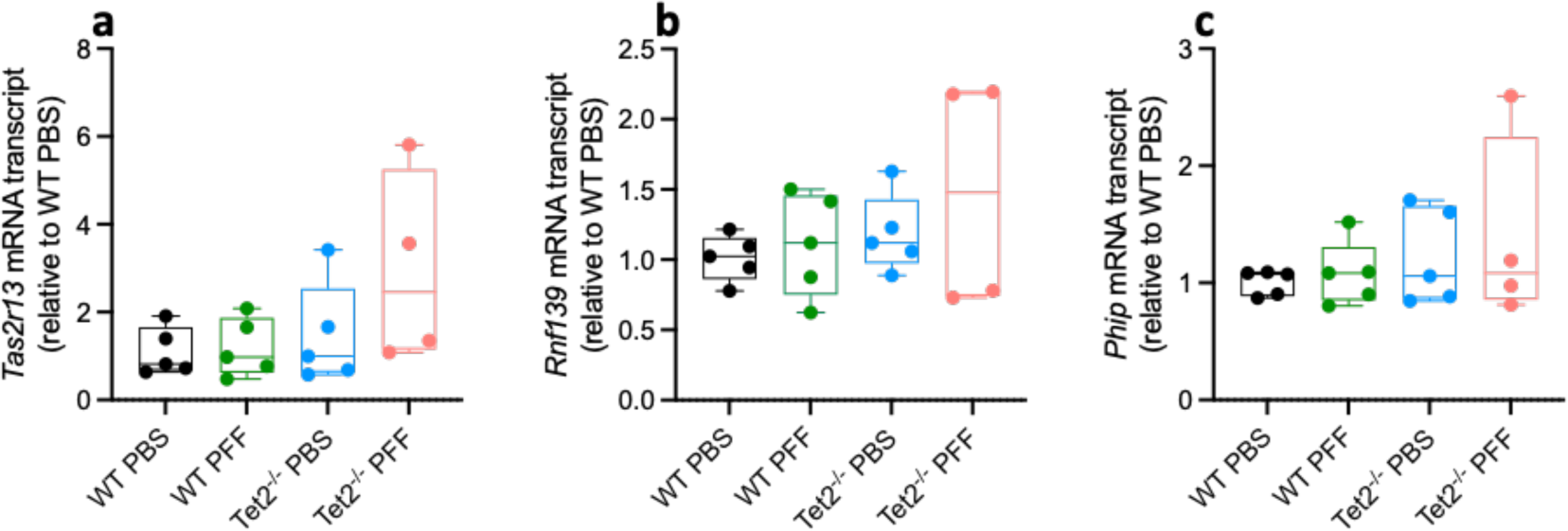
*Tet2* deletion does not alter the expression of selected genes dysregulated in PD neurons. Six months after striatal αSyn PFF inoculation in WT and *Tet2*-knockout mice, we examined cortical expression of three randomly selected genes that are dysregulated in PD neurons, **a** *Tas2r13,* **b** *Rnf139*, and **c** *Phip* (*n*=4-5 mice per group). Data are mean ± s.e.m.

**Supplementary Fig. 6.**
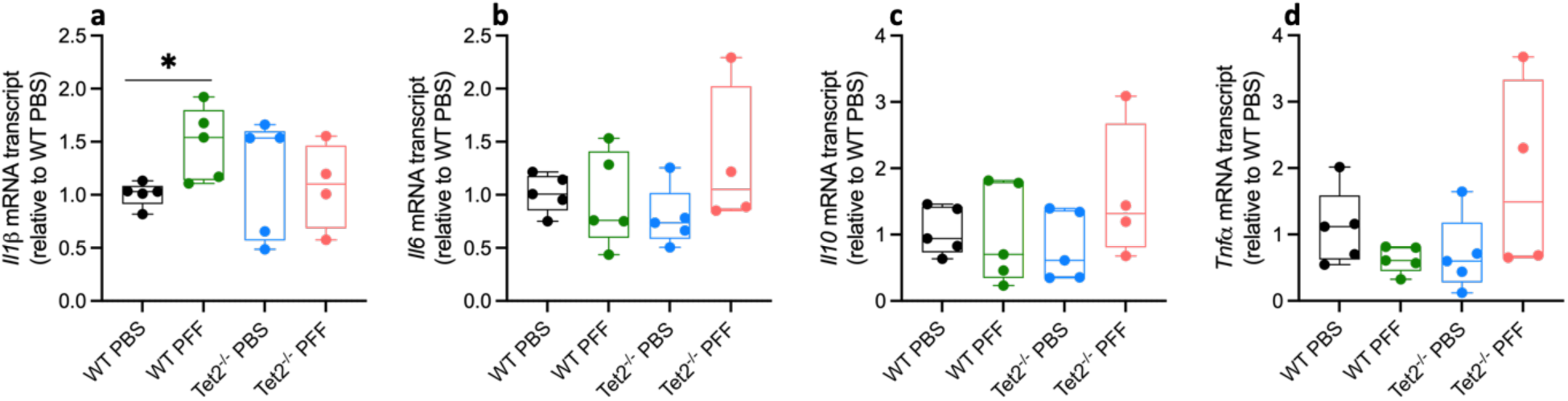
*Tet2* deletion normalizes the increased expression of *il1β* induced by αSyn pathology. Six months after striatal αSyn PFF inoculation, we examined cortical cytokine expression via qPCR, **a** mRNA expression of *il1β*, **b** *il6*, **c** *il10,* and **d** *Tnfα* (*n*=4-5 mice per group). Data are mean ± s.e.m., analyzed using two-sided t-test, \**P* < 0.05.

**Supplementary Table 1** Demographic and clinical information for study samples.

**Supplementary Table 2** Primers used in detecting the expression levels of target genes.

